# *In situ* novel environment assay reveals acoustic exploration as a repeatable behavioral response in migratory bats

**DOI:** 10.1101/2020.12.16.423043

**Authors:** Theresa Schabacker, Oliver Lindecke, Sofia Rizzi, Lara Marggraf, Gunārs Pētersons, Christian C. Voigt, Lysanne Snijders

## Abstract

Integrating information on species-specific sensory perception together with spatial activity provides a high-resolution understanding of how animals explore environments, yet frequently used exploration assays commonly ignore sensory acquisition as a measure for exploration. Echolocation is an active sensing system used by hundreds of mammal species, primarily bats. As echolocation call activity can be reliably quantified, bats present an excellent animal model to investigate intra-specific variation in environmental cue sampling. Here, we developed an *in situ* roost-like novel environment assay for tree-cave roosting bats. We repeatedly tested 52 individuals of the migratory bat species, *Pipistrellus nathusii*, across 24 hours, to examine the role of echolocation when crawling through a maze-type arena and test for consistent intra-specific variation in sensory-based exploration. We reveal a strong correlation between echolocation call activity and spatial activity. Moreover, we show that during the exploration of the maze, individuals consistently differed in spatial activity as well as echolocation call activity given their spatial activity, a behavioral response we term ‘acoustic exploration’. Acoustic exploration was correlated with other exploratory behaviors, but not with emergence latency. We here present a relevant new measure for exploration behavior and provide evidence for consistent (short-term) intra-specific variation in the level at which wild bats collect information from a novel environment.

## Introduction

Sensory systems are fundamental to survival and reproduction, acting at the interface between an organism’s central nervous system and its environment^1,2^. Sensory systems allow organisms to collect, process and respond adequately to environmental information, thereby reducing uncertainties about aspects of their environment^3^. Animals make decisions that are strongly influenced by the information they have of their environment^4^, which in turn is based on the information that was transferred through their sensory systems^2,5^. Sensory systems can include ‘active’ and ‘passive’ sensing^6^. Passive sensing (*e.g*. vision) relies on sources that generate or reflect energy (*e.g*. electromagnetic energy from the sun), extrinsic to the sensing organism. Active sensing relies on the use of self-generated energy to receive information from the environment (*e.g*. echolocation), enabling animals to control characteristics of the probe energy, such as intensity, direction and timing^6^.

The collection of environmental cues, stemming from active or passive sensing, is reflected by a behavior commonly referred to as ‘exploration’^7^. Through exploration animals gain vital information about the availability of resources, the distribution of conspecifics, and the presence of predators. Animals frequently get confronted with changes to their environment, which have the potential to impact an individual’s fitness^8,9^. Currently, anthropogenic change, and particularly climate change, is impacting the distribution and presence of mates, predators, prey and suitable habitat in both space and time^10–13^. Gathering detailed and up-to-date environmental information via exploration can help animals anticipate and mitigate such disruptive changes to their environment.

Former studies often assessed exploration levels in wild animals by measuring spatial activity as a proxy for exploration, by considering the speed of exploration in a novel environment or the proportion of a new environment explored (as discussed by^14,15^). Interestingly, many of these investigations revealed consistent differences in exploration levels among individuals of the same species (reviewed in^16,17^), which were also found to be heritable in some species, such as the great tit, *Parus major*^18,19^. Individual differences in exploration are not trivial^17^, as they may correlate with foraging strategies^20,21^, dispersal^22^ or competitive ability^23^. Yet, exploration, when defined purely by spatial measures, may primarily represent activity-level and not necessarily reflect the number or quality of environmental cues being collected by the individual^14^, hampering our mechanistic understanding of how intra-specific variation in exploration and environmental change are likely to interact and shape natural selection of a species.

Measuring exploration, including a measure for environmental cue sampling, is challenging in species which primarily use passive sensing, such as birds, (but see^15^). Bats, however, are a suitable study model to circumvent these limitations. Except for members of the family of Pteroporidae, bats have adapted to their nocturnal niche by using laryngeal echolocation^24^. Echolocation is an active acoustic sensing system, enabling bats to individually control the incoming information about the environment, by actively manipulating frequency, direction, and timing of echolocation pulses. Modern technology now allows us to reliably quantify echolocation calls in the field. Echolocation call activity thus provides us with the exciting opportunity to investigate intra-specific variation in how animals sample cues when exploring novel environments.

Bats use echolocation to orchestrate a variety of tasks, starting with the assessment of environments (*i.e*. orientation) in the complete darkness of their roosts^25^ and under nightly low light conditions in flight. Echolocation supports hunting of prey on the wing^26^, communication^27^ and navigation during wayfinding using landmarks^28,29^. The Nathusius’ pipistrelle (*Pipistrellus nathusii*), a tree-cave roosting insectivorous bat, is widely distributed in Europe and migrates every year between breeding grounds in northeastern Europe and hibernation areas in southwestern Europe^30–33^. Migratory bats are particularly vulnerable to environmental changes as they depend on specific conditions on a global geographical scale rather than in just one specific habitat^34^. Moreover, Nathusius’ pipistrelles suffer from extensive mortalities at wind turbines^35,36^. For constructive conservation plans, it is therefore necessary to gain deeper understanding of how bats perceive and react to environmental cues and to uncover potential variation among individuals.

Here, we developed an emergence and novel environment assay for tree-cave roosting bats with laryngeal echolocation to be used in the field. At a migratory corridor at the coastline of Latvia, we repeatedly assayed (24 hour time-interval) 52 male and female Nathusius’ pipistrelles, quantifying behavior for the first two minutes following emergence into a roost-like novel environment, consisting of a maze-type arena with nine chambers through which the bat could crawl. We (1) tested whether echolocation call activity related to spatial activity in the novel environment. If echolocation call activity is relevant to exploration, we would expect the acoustic and spatial responses to be positively correlated. To examine whether echolocation call activity provided information about exploration behavior in addition to spatial activity, we quantified the number of echolocation calls while controlling for spatial activity and (2) tested whether this ‘acoustic exploration’ behavior was individually repeatable, *i.e*. whether certain individuals consistently under- or oversampled a novel environment. Lastly, we (3) tested whether acoustic exploration was related to emergence latency, as proxy for boldness, to evaluate the potential presence of a behavioral syndrome^37^.

## Results

### Variation in behavioral responses while exploring a novel environment

Bats fully entered the maze in 91 out of 111 assays (82%), representing 59 unique individuals (Fig. 1a). In addition, four bats emerged with their head but never with their full body. Fully emerged bats showed substantial variation in all analyzed behavioral responses (Supplementary Table 1), including emergence behavior, *e.g*. latency of body emergence (1 to 162 seconds, Fig. 1a) and duration between head and full body emergence (1 to 114 seconds, Fig. 1c), acoustic behavior, *e.g*. number of echolocation calls during emergence (4 to 335, Fig. 1d) and number of echolocation calls during the first two minutes after emergence (170 to 2127, Fig. 1e) and spatial behavior, *e.g*. number of unique chambers (1 to 9, Fig. 1f) and total number of chambers (1 to 25, Fig. 1g) visited during the first two minutes in the novel environment. Furthermore, the number of times a bat poked its head in an adjacent chamber ranged from 0 to 7, the number of emitted ‘air puffs’, *i.e*. brief vocal outbursts audible to the human ear, ranged from 0 to 22 and in 23 out of 91 assays a bat (also) crawled through the novel environment upside down (Supplementary Table 1).

**Figure 1.**
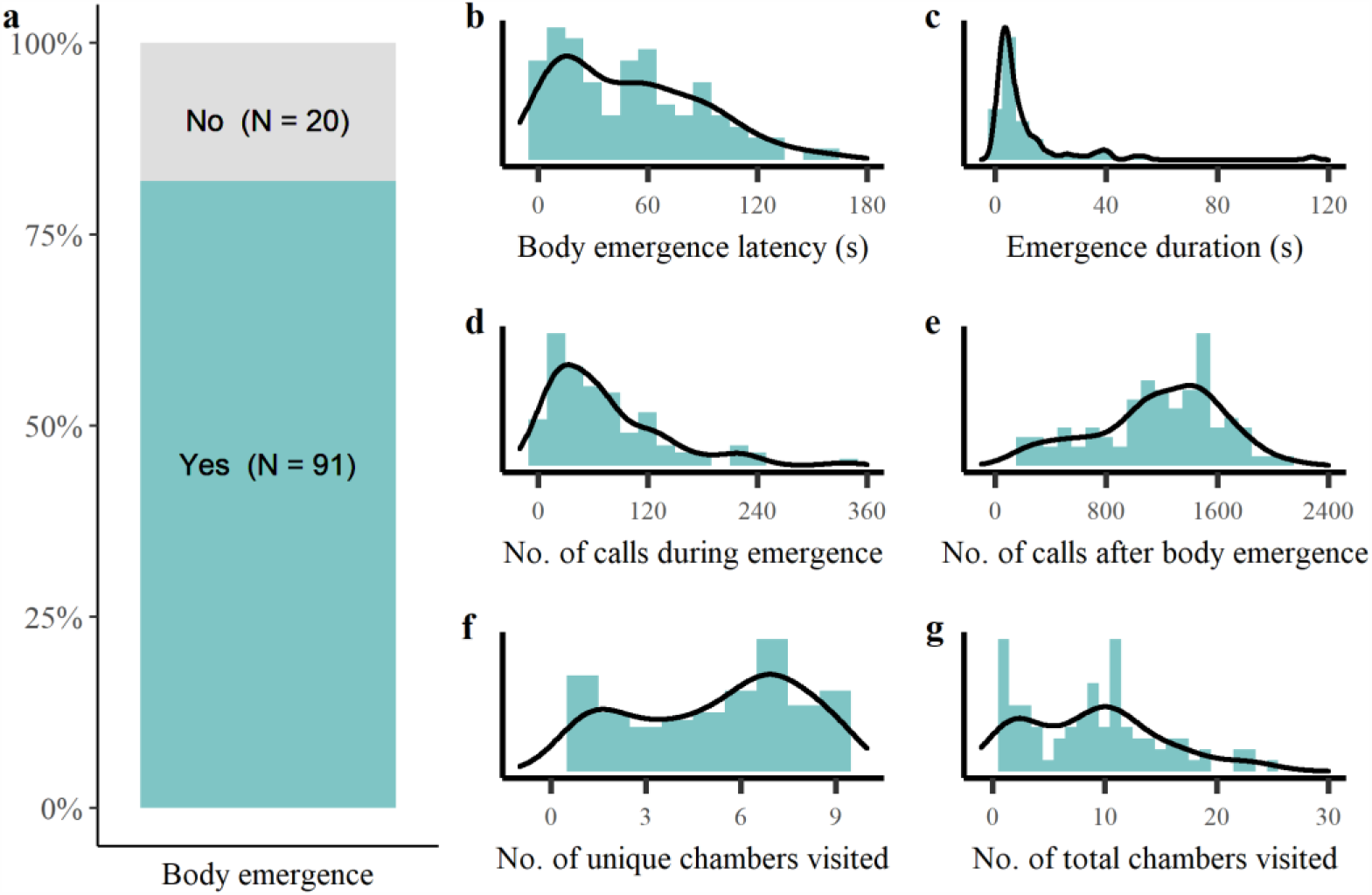
Variance in behavioral responses in Nathusisus’ pipistrelles exploring a novel environment. (a) In 82% of the tests the bat fully emerged out of the starting tube and entered the novel environment (a maze-like arena). The substantial variation in behavioral response to the novel environment is visualized for (b) latency to enter the novel environment (s), (c) duration between head and full body emergence (s), (d) number of echolocation calls during emergence (*i.e*. time between head and body emergence), (e) number of calls after full body emergence, (f) number of unique chambers visited and (g) total number of chambers visited during the first two minutes in the novel environment. The lines in panels b-g show the density distribution while the bars show the frequency distribution.

### Echolocation call activity positively correlates to spatial activity in a novel environment

Different response measures of the same behavioral category were moderately to strongly correlated to each other (r_s_ = 0.25 -0.93 for the emergence measures, r_s_ = 0.22 - 0.98 for the acoustic measures and r_s_ = 0.89 for the spatial measures, Supplementary Table 2). The acoustic and spatial response to a novel environment were strongly correlated to each other, with bats that visited more chambers also emitting more echolocation calls (r = 0.65, p < 0.001, Fig. 2a). Nevertheless, there was still substantial variation among bats visiting a similar number of chambers (Fig. 2a), a behavioral response we termed ‘acoustic exploration’ (Fig. 2b).

**Figure 2.**
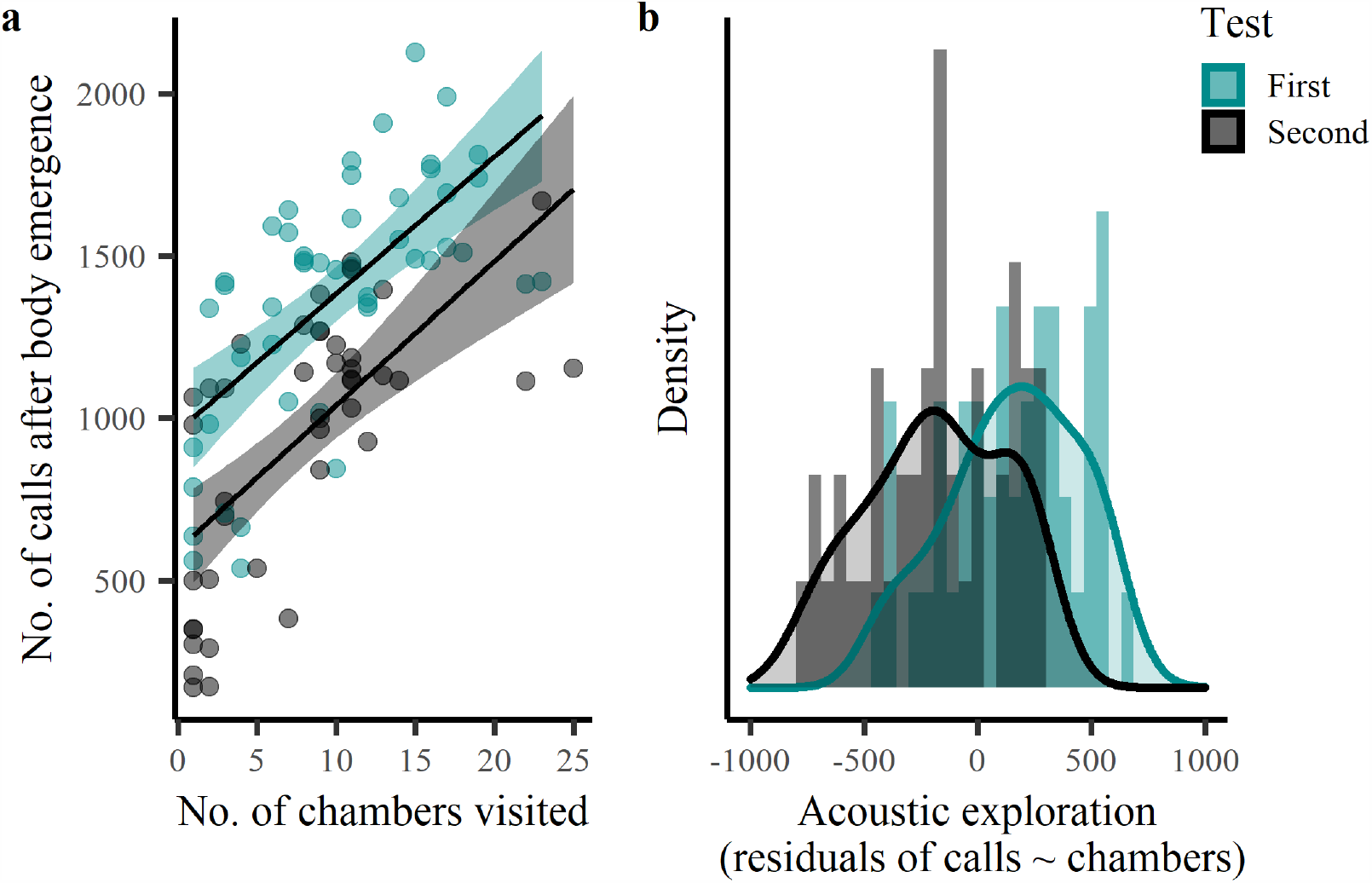
Number of echolocation calls emitted during the first and second test as function of the number of chambers visited by Nathusisus’ pipistrelles in a novel environment. (a) Bats emitted more echolocation calls during the first two minutes in the novel environment (a maze-like arena) when they also visited more chambers. (b) Bats varied substantially in how much more or less they echolocated than expected based on the number of chambers they visited, *i.e*. the residuals of the number of echolocation calls as function of the number of chambers visited varied considerably. Regression lines in panel (a) show the predicted final model values. Shaded areas around the lines reflect the 95% Confidence Interval. Lines in panel (b) show the density distribution while the bars show the frequency distribution.

### Individuals consistently under or oversample a novel environment

We were able to conduct two valid novel environment assays (approximately 24 hours apart) with 52 unique individuals. Bats emerged in 85 out of these 104 tests (82%). Acoustic exploration was significantly reduced during the second assay (Estimate (Est.) ± Standard Error (SE): -344.60 ± 52.43, p < 0.001, Fig. 3b, Supplementary Table 3) and males showed more acoustic exploration than females (Est. ± SE = 144.71 ± 69.44, p = 0.04, Fig. 3c, Supplementary Table 3). Bats showed more acoustic exploration when the set-up faced more South compared to North and more East compared to West (North-South: Est. ± SE = -93.99 ± 43.81, p = 0.03; East-West: Est. ± SE = 85.95 ± 42.36, p = 0.04, Supplementary Fig. 1, Supplementary Table 3). Interestingly, individuals consistently differed in how thoroughly they acoustically explored the novel environment, i.e. the relative number of echolocation calls they emitted per chamber (R_adj_ = 0.32, SE = 0.15, CI = 0.07 - 0.62, p = 0.03, Fig 3a).

**Figure 3.**
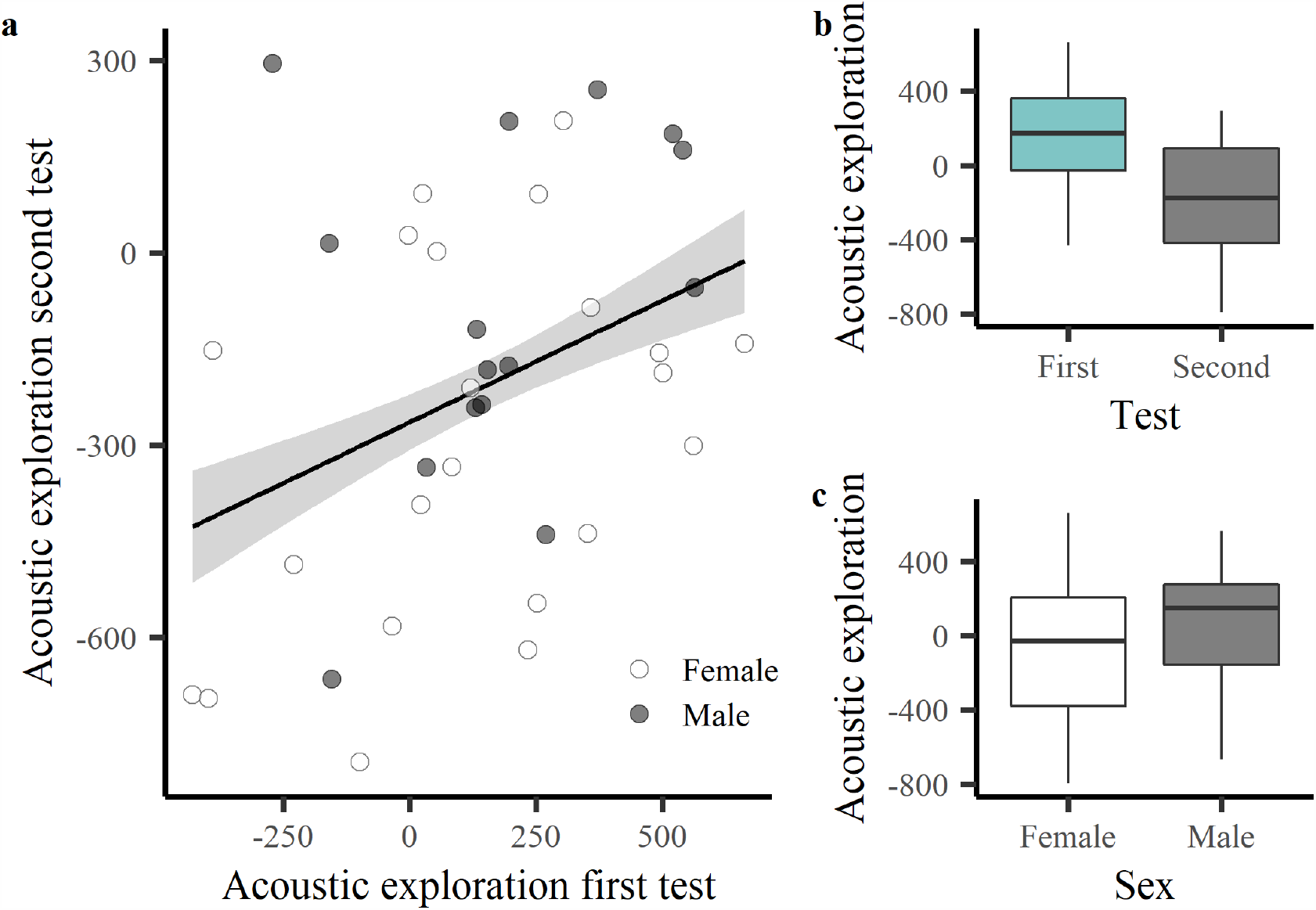
Acoustic exploration, *i.e*. residuals of the number of echolocation calls as function of the number of chambers visited, during the first and second novel environment test with Nathusius’ pipistrelles. (a) Acoustic exploration during the first novel environment test (a maze-like arena) predicts an individual’s acoustic exploration during the second test. (b) Individuals show more acoustic exploration during the first than the second test. (c) Males show more acoustic exploration than females. Males and females are represented by a black or white fill, respectively. The regression line in panel (a) shows the predicted final model values. Shaded areas around the line reflects the 95% Confidence Interval. Box plots in panel (b) and (c) show median and interquartile range with whiskers of 1.5 interquartile distances.

Bats also tended to visit fewer chambers during the second test (Est. ± SE = -0.28 ± 0.14, p = 0.053, Supplementary Table 4) and individuals consistently differed in how many chambers they visited (R_adj_= 0.29, SE = 0.16, CI = 0 - 0.58, p = 0.048). Test number did not significantly influence emergence latency (Est. ± SE = -0.34 ± 0.23, p = 0.13, Supplementary Table 5) and individuals that emerged quicker during the first test tended to also emerge quicker during the second test (R_adj_ = 0.23, SE = 0.13, CI = 0 - 0.50, p = 0.054). Yet, it should be noted that no repeatable individual differences in emergence latency could be detected when the minimum latencies for non-emergers were completely excluded (181 s, 18% of tests).

### Acoustic exploration correlates to other exploratory behaviors but not to emergence latency

Bats that emerged quicker (or slower) did not exhibit more acoustic exploration (Est. ± SE = 5.00 ± 27.48, p = 0.86, Fig. 4a, Supplementary Table 6). However, bats that emitted relatively more echolocation calls during emergence (Residuals of: Number of calls during emergence ∼ Emergence duration (s): r_s_ = 0.79, p < 0.001) similarly showed more acoustic exploration, *i.e*. after emergence (Est. ± SE = 2.36 ± 0.57, p < 0.001, Fig. 4b, Supplementary Table 6). In addition, bats that poked their heads into an adjacent chamber more often also showed more acoustic exploration (Est. ± SE = 57.87 ± 19.16, p = 0.003, Fig. 4c, Supplementary Table 6). The number of air puffs a bat emitted did not correlate with acoustic exploration (Est. ± SE = 7.71 ± 7.85, p = 0.33, Supplementary Table 6). Finally, there was a trend for bats that crawled upside down for part of the test to exhibit less acoustic exploration (Est. ± SE = -128.15 ± 68.94, p = 0.06, Supplementary Table 6).

**Figure 4.**
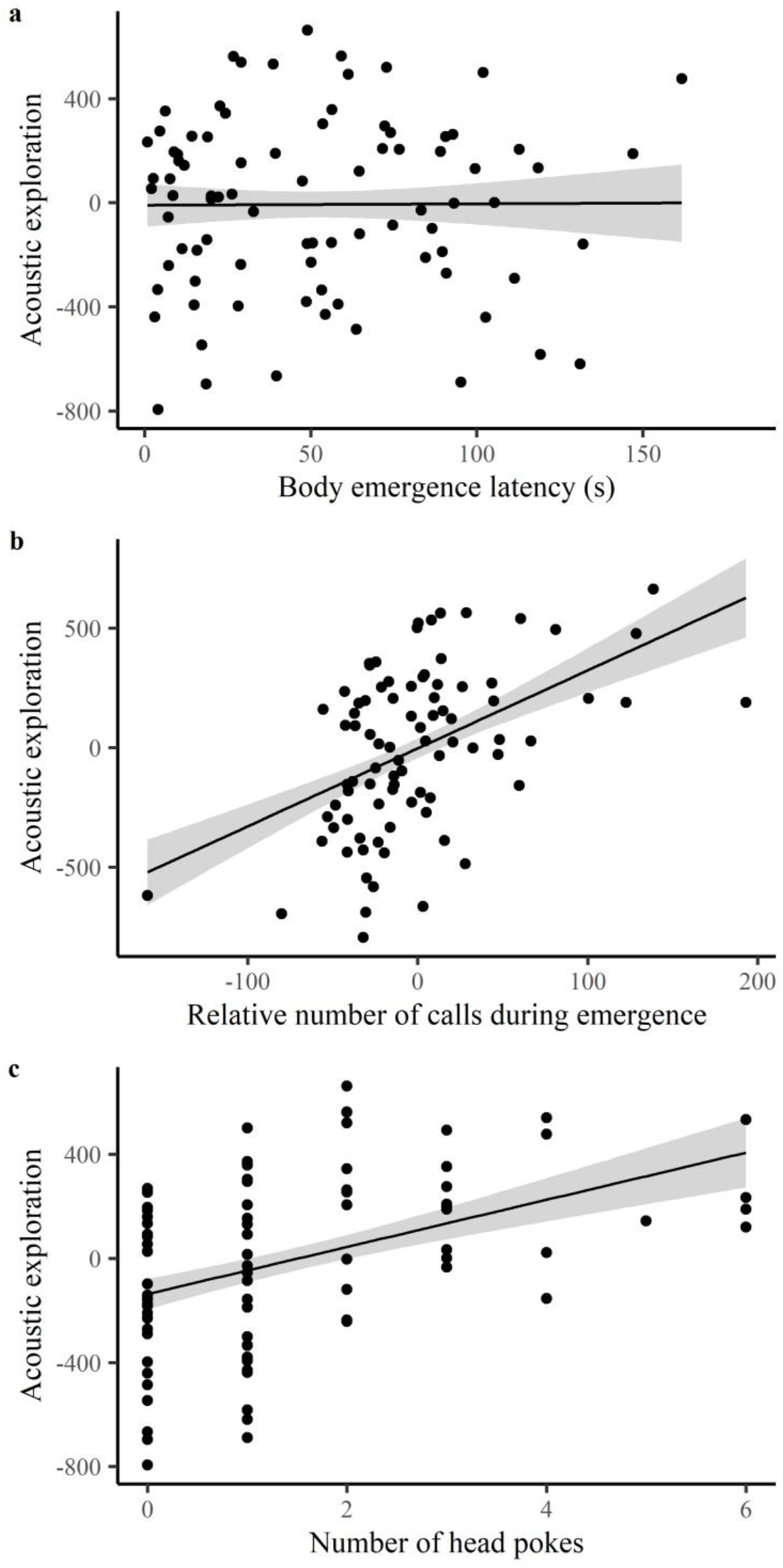
Acoustic exploration, *i.e*. residuals of the number of echolocation calls as function of the number of chambers visited, in relation to other behavioral responses exhibited by Nathusius’ pipistrelles in a novel environment. These behavioral responses exhibited in the novel environment (a maze-like arena) included: (a) Body emergence latency (s), (b) Acoustic exploration during emergence, i.e. the residuals of the number of echolocation calls as function of emergence duration (time between the head out and the body out of the starting tube), (c) the number of times a bat poked its head into an adjacent chamber without entering. Regression lines show the predicted final model values. Shaded areas around the lines reflect the 95% Confidence Interval.

## Discussion

Animals use their sensory systems to explore (*i.e*. examine and investigate) environments and objects, a behavior crucial for animals to keep up to date with the presence and distribution of resources and threats and so reduce uncertainty about their environment^38^. Yet, most of our current understanding of variation in exploration behavior is based on where animals go rather than the information they collect. Here, we reveal a relevant new measure for exploration behavior by quantifying sensory activity relative to spatial activity and subsequently providing evidence for consistent (short-term) variation between individual bats in how thoroughly they acoustically explore a novel environment.

With our newly developed *in situ* assay, we were not only able to uncover substantial intra-specific variation in bats’ responses to a novel environment, but also to reveal that individual bats consistently varied in these responses. Spatial movement and acoustic activity, both highly relevant behaviors in the context of orientation and navigation^28,39^, were strongly positively correlated. In line with previous studies investigating exploration behavior in other species (*e.g*. fish^21,40^, amphibians^41^, birds^42,43^, mammals^44,45^), we found spatial activity in a novel experimental arena to be individually repeatable. We add to these studies by now showing that also sensory activity (*e.g*. number of echolocation calls), given spatial activity, is an individually repeatable response, which we termed ‘acoustic exploration’. Our repeatability estimate for acoustic exploration (R = 0.32) was similar to those reported for a range of behavioral traits across a wide variety of animal taxa (average R = 0.37)^16^. Importantly, acoustic exploration correlated with the number of times an individual poked its head into an adjacent chamber and the relative echolocation call rate during emergence, suggesting acoustic exploration to represent true exploration behavior, rather than activity. Exploration behavior frequently correlates with other behavioral traits, including aggressiveness^46^ and boldness^47^, forming a behavioral syndrome^37^. Though, we did not find evidence for a correlation between acoustic exploration and emergence latency, an often-used proxy for boldness (*e.g*.^47^), in this study. Recent examination of *in situ* takeoff orientation in this species using a circular release box set-up^48^ revealed emergence latency values that were highly similar (range: 6 – 176 s, median: 55 s) to those we report here (Supplementary Table 1). Both set-ups confronted individuals with a multiple directional choice for movement in a novel environment, raising the possibility that not only risk-aversion but also decision or orientation-speed may have influenced emergence latency, a hypothesis that requires further testing.

Interestingly, exploration behavior has also been linked to routinization^49^ and learning ability^43^. Thorough exploration likely creates a greater knowledge of the state and quality of the habitat, allowing individuals to better react to spontaneously arising environmental stimuli^50^. Building forward on this, researchers have proposed the existence of a ‘cognitive syndrome’, based on the assumption that individuals differ in the speed and accuracy of picking up environmental cues leading to two alternative cognitive styles, namely ‘fast and superficial’ and ‘slow but accurate’^51^. This theoretical framework suggests that some individuals initially learn about specific contingencies more quickly, potentially by recognizing and memorizing them faster and by being more susceptible to forming behavioral routines^52^. However, as the formation of behavioral routines is associated with some degree of inflexibility, these individuals are expected to be more challenged in tasks that require behavioral alterations in response to altered environmental conditions^53,54^. Thorough exploring bats may thus have an evolutionary advantage when facing novel threats and habitat structures (*e.g*.^55,56^), as an increase in sensory update rate is thought to allow faster behavioral responses^39,57^. Thorough explorers can thus be expected to detect changes in the environment more quickly^58^, increasing their adaptive potential and resilience to environmental change^59,60^. For example, exploration behavior in a novel environment by Iberian wall lizards’ (*Podarcis hispanica*) was positively associated with the habituation speed to a threatening stimulus^61^. Linking individual variation in acoustic exploration to the ability of bats to quickly detect and respond to novelty on a wider spatial scale, or a comparative investigation of bat populations living along a gradient of habitat stability, would be exciting next steps. Furthermore, a more intense use of an active sensory system, such as echolocation, is not necessarily without energetic costs^62^, so an examination of the potential costs linked to more thorough exploration in bats would likewise be insightful.

Our behavioral repeatability estimates were based on a short time-interval (24 hours) and ideally longer intervals would be used. However, it is logistically extremely challenging to assess wild migratory and nocturnal animals over a longer time span. We therefore see our study as a first step towards evidence for consistent individual differences in acoustic exploration behavior in wild migratory bats and encourage follow-up studies employing longer timespans to further investigate the stability of these differences. We also witnessed reduced acoustic exploration and spatial activity during the second assay which were likely caused by habituation. Habituation is a common problem in novelty tests as repeated exposure inherently reduces novelty and creates a sense of familiarity^63,64^. As a result, the motivation, or the need, to explore diminishes as the environment becomes more familiar^65–67^. However, habituation also has adaptive significance, as it allows animals to separate repeated, potentially irrelevant, stimuli from important ones^68^. Even though we were still able to reveal individual repeatability in exploration behavior, future studies could possibly control for habituation effects by deploying a different type of novel environment during the second assay. Such a set-up could also be adapted to investigate the presence of consistent individual differences in habituation, by repeatedly testing habituation to novelty, using a reaction-norm approach^69^.

Despite their high diversity and multitude of ecosystem services, few studies have investigated consistency of intra-specific variation in bats and how this variation may influence a species fitness and ecology. Some exceptions include studies on species from the family of Vespertilionidae that showed these bats to exhibit consistent intra-specific variation in (aggregate) behavioral responses linked to exploration, activity, anxiety^70^ and aggressiveness^71^. Moreover, this variation was linked to pathogen dynamics^72^. Another major selection process in the lives of many bat species, is habitat quality. For tree-dwelling bat species, roosts are crucial for resting^73^, reproducing^74^ and mitigating predation risk^75^, thus roost deficits could be detrimental for bats^76^. However, the large majority (97%) of European forests are managed and under commercial use^77^, *e.g*. felling of trees and removal of deadwood, resulting in a reduced habitat quality for bats^78^. Especially for migrants, knowledge about adequate roosts along the route is vital and bats can be expected to remember roost locations even over hibernation^79^. If trees or forest areas with suitable roosts are lost due to anthropogenic activities, these conditions may favor high levels of exploration and/or plasticity. Thus, thorough explorers may have a selective advantage, resulting in a population-level shift in exploration behavior and associated behaviors and a potential increase in intra-specific competition^80^.

For the Nathusius’ pipistrelle, the time of late summer to autumn migration overlaps with the time of mating, which peaks in August in Latvia^30,32,81^. During this period, males are typically more conspicuous, as they produce loud acoustic displays as advertisement calls during flight or while roosting, which are believed to repel other males and/or attracts females^82^. This may partially explain the higher acoustic exploration of males in this study. The observed higher levels of acoustic exploration when the starting tube of the maze was facing South or East may be related to bats’ migratory behavior. Nathusius’ pipistrelles appear to have an intrinsic orientation, as these bats showed consistent departure flight directions in the migratory direction, even when translocated more than 10 km away from capture site^83^. It is therefore possible that the alignment of the maze with the migratory direction stimulated enhanced acoustic exploration, but further hypothesis-based testing would be necessary to draw reliable conclusions.

## Conclusion

Evaluating animal exploration is a challenging task, as this behavior is defined in terms of information acquisition^7^, but classically measured in terms of movement speed or proportion of a novel environment explored^15,84^. Here, we address this challenge by developing an *in situ* assay to assess echolocation call activity relative to spatial activity while exploring a novel environment. The assessment of acoustic exploration in a tree-cave roosting bat species proved to be a very suitable approach to obtain high-resolution sensory-based data on individual behavior in response to a novel environment. Next to providing novel evidence for an exploration-relevant repeatable behavioral trait, we hope our approach instigates further research into intra-specific variation in environmental cue sampling, using species with predominant use of active sensing modalities as study models, and inspire a species-specific sensory perspective on how individual animals may respond differently to environmental novelty.

## Materials and Methods

### Study population

60 adult migratory Nathusius’ bats were caught at the Pape Ornithological Station, University of Latvia, Latvia (56°09′N, 21°03′E) between 09:00 PM and 03:00 AM on the 16^th^ and 18^th^ of August 2019, using a Helgoland funnel trap^32^. The study adhered to ASAB/ABS guidelines for the treatment of animals. The general set-up and protocol for assessing exploration behavior in tree-cave roosting bats was evaluated by the animal welfare representatives of the Leibniz Institute for Zoo and Wildlife Research. The work was conducted under permit no. 49/2019 issued by the Latvian Nature Conservation Agency to G.P.

### General procedure

Following capture, we determined the sex, body mass (digital balance) and forearm length (manual calipers). Bats received a temporary ID coded gummy ring for individual identification. The bats were kept together in a dark and quiet environment (dark, wooden shed) in wooden carriers with a mesh top and covered by a dark cloth (seven to eight individuals per 30 cm x 30 cm x 10 cm), mimicking natural day-time roost conditions during which bats save energy by going into torpor. The bats general condition and body mass was checked daily, and water and food were offered, with individuals weighing less than seven grams being given supplemental feeding (Harrison’s Juvenile Hand-Feeding Formula). Behavioral assays were conducted on two consecutive nights following capture and took place in a dark wooden cabin (3 m x 2 m, constant temperature of 21 ± 2 °C) within 200 m of the location of capture.

### Set-up and protocol

Prior to an assay, a bat was kept in an individual cotton bag (Ecotone, Gdansk, Poland) near a wrapped-up warm water bottle to stimulate it to come out of torpor. The bat’s normothermia was confirmed by measuring the skin temperature (> 30° C) with a thermocouple (Peakmeter, PM6501; Thermocouple, Sensor SSP-1-150, Peakmeter, Shenzhen, China). Skin temperature has been shown to be a non-invasive measurement that accurately reflects rectal temperature^85^.

We encouraged natural exploration behavior by constructing an experimental maze-like arena, mimicking a novel environment that would be relevant for a tree-cave roosting bat species (Figure 5; 40 cm × 40 cm × 5 cm). The maze, placed in a larger box (70 cm x 45 cm x 8 cm) with a transparent plastic lid, consisted of nine separate chambers accessible through small gates (3 cm × 2.5 cm) on the upper half of the walls (Fig. 5). The maze was placed horizontally to stimulate exploratory movements in all directions. The floor was covered with a plastic non-slip rug mat to supply the bat with a good texture for crawling, a common behavior for tree-cave roosting bats. A layer of insect screen covered the maze (not shown in Fig. 5), offering additional climbing opportunities and preventing the bat from escaping. This set up enabled us to capture the bat’s movement via a night vision camera (Sony Digital Camcorder, DCR-SR72E, Sony, Tokyo, Japan) mounted on a tripod with a horizontal arm to allow an aerial view from 1.5 m. An infra-red torch (T38, Evolva Future Technology, Shenzhen, China), shining at an angle from a fixed position, provided light for the camera (the camera’s own light was taped off to prevent reflection). A microphone (USG Electret Ultrasound Microphone, Avisoft Bioacoustics, Berlin, Germany) connected to an ultrasound recording device (UltraSoundGate 116Hb, Avisoft Bioacoustics, Berlin, Germany) was placed within the larger box to record all vocalizations. An opaque start tube (10 cm × 3 cm) was attached to the maze but blocked by a small wooden barrier which we removed at the start of the assay (Fig. 5).

**Figure 5.**
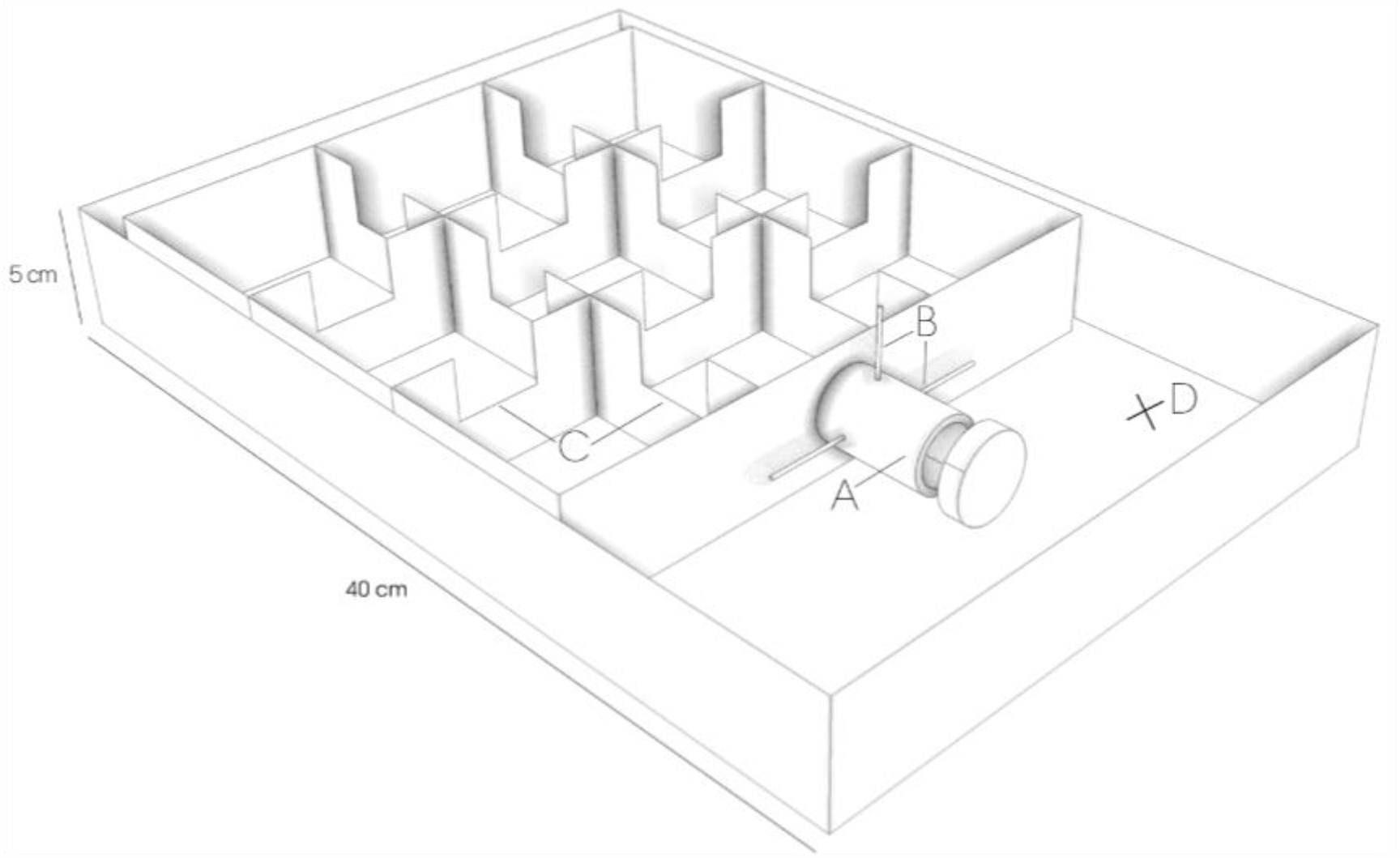
Schematic drawing of the maze used during behavioral assay. A) Opaque start tube where bats were placed at the start of each assay B) Barriers closing entrance to maze C) Gates connecting single chambers D) Position of microphone. ©Rebecca Scheibke

We placed the bat in the starting tube, with the exit obstructed by vertical and horizontal barriers in the form of wooden dowels. After 20 seconds we removed the vertical barrier and waited a maximum of three minutes for the bat to emerge^48^. The time taken for the bats to completely emerge from the start tube was recorded. If the bat did not emerge within this time, the test was terminated. After emergence, bats were given two minutes to explore the box. After each test, we sterilized the maze with a mild unscented detergent to remove potential olfactory cues left by previous subjects. Halfway through the night we rotated the assay to control for potential orientation biases. Flight orientation was briefly assessed on the same night (different study) and controlled for in the analysis (see ‘Statistical analysis’). Bats were released the night of the second assay within 100 m of the original capture location.

### Behavioral response quantification

We used the event-logger BORIS^86^ version 7.9.1 to quantify several behavioral response measures in detail: latency to head emergence (s), latency to full body emergence (s), number of unique chambers discovered, total number of chambers visited, number of times a bat poked its head in an adjacent chamber and whether a bat had crawled upside down through the maze. In addition, we recorded the echolocation activity during emergence as well as after head and full body emergence and quantified the number of echolocation calls using Avisoft SASLab Pro version 5.2 (©Avisoft Bioacoustics, Berlin, Germany). Prior to analysis, sampling frequency was converted from 250 kHz to 150 kHz. The spectrogram was computed with Fast Fourier Transformation 256, parameters were set to Hamming window (bandwidth 1270 Hz, resolution 977 Hz) and volume normalized to 75%. We synchronized the audio recordings with the video recordings via spoken start and stop commands given by the observer. We used Avisoft’s call detection and template-based spectrogram comparison feature to identify distinct echolocation calls, which were verified and corrected via visual inspection (correlation corrected and uncorrected: rs > 0.99, p < 0.001). Finally, we quantified the number of ‘air puffs’, brief audible acoustic outbursts that sound similar to sneezes.

### Statistical analysis

We conducted all analyses with R^87^ version 4.0.2 in R Studio version 1.2.5033 (© 2009–2019 RStudio, Inc.). All statistical tests were two-sided. We first conducted Pearson (parametric) and Spearman (non-parametric) correlation tests between measures within a behavioral category (*e.g*. acoustic behavior), using all valid test data (n = 111 tests without premature emergence, 59 individuals), to evaluate whether we could proceed with the analysis using one representative response measure and so minimize redundancy. Next, using a Pearson correlation test, we tested whether the number of echolocation calls following the first two minutes since emergence correlated to the number of chambers visited. Subsequently, we extracted the residuals for this correlation (*i.e*. the ‘acoustic exploration’ measure) using a linear model. We additionally tested whether the number of echolocation calls emitted during emergence was correlated to the duration of emergence (seconds between head and full body emergence) using a Spearman correlation test and extracted the residuals using a linear model.

Three mixed model selection procedures were conducted using the lmer and glmer functions of the ‘lme4’ package^88^, with the aim of identifying variables that significantly affect (1) acoustic exploration, (2) number of chambers visited and (3) emergence latency. These models only included data for the 52 individual bats (n_♀_ = 31, n_♂_ = 21) for which we had two valid test responses. We used stepwise backward model selection, assessing the significance of fixed effects by the change in deviance upon removal of the effect (Type II Wald Chi-square tests). Effects with p < 0.1 were kept in the final model. All models included individual ID as a random effect and all three starting models included the independent variables: test (two-level factor), sex (two-level factor), weight in grams (scaled covariate), forearm length in mm (scaled covariate), time since sunset (scaled covariate), direction of starting tube (two integer variables: North-South with North = 1, South = -1, East and West = 0; East-West with East = 1, West = -1, North and South = 0), supplemental food (two-level factor) and orientation assessment (two-level factor). The factor ‘test’ was kept in the model at all times. Potential collinearity was assessed using the variation inflation factor (VIF) extracted with the ‘performance’ package^89^ and variables above 2.5 are reported in the Supplementary Information (none of our variables of interest, but some of our control variables showed collinearity).

First, effects on acoustic exploration were analyzed using a linear mixed model (LMM). The model assumptions were evaluated using the Shapiro-Wilk test and a visual inspection of the model residuals. Second, effects on the visited number of chambers were analyzed using a generalized linear mixed model (GLMM) with a poisson error-distribution and the bobyqa optimizer. Following overdispersion, an observation-level random effect (*i.e*. observation number) was added to the model. Third, effects on body emergence latency were analyzed by log-transforming emergence latency and using an LMM. Values for non-emergers were set to the minimal possible value of 181 seconds. Although the Shapiro-Wilk test indicated a significant deviation from normality during the model selection steps for emergence latency (0.02 < p < 0.03), the histogram of the residual frequency distribution did not show substantial deviations and the check_distribution function of the ‘performance’ package indicated a 100% probability of the model residuals originating from a normal distribution.

Repeatability (R_adj_) of acoustic exploration, number of visited chambers and emergence latency (log-transformed) was quantified with the ‘rptR’ package^90^ using the final model from the model selection procedure. Confidence intervals for the repeatability estimates were bootstrapped (1000 times). To evaluate potential correlations between acoustic exploration and other behavioral response measures, the respective response measure was added to the final model for acoustic exploration and significance was assessed using Type II Wald Chi-square tests. Figures were created using the ‘ggplot2’ package^91^.

## Acknowledgements

We are grateful to the many fieldworkers of the Pape Ornithological Station. We thank Shannon Currie for assistance and expert advice during the fieldwork. L.S. was funded by a Humboldt Research Fellowship for Postdoctoral Researchers (Ref 3.3 - NLD - 1192631 - HFST-P) awarded by the Alexander von Humboldt-Stiftung. We thank Rebecca Scheibke for her illustration of the novel environment assay.

## Author contributions

L.S., O.L, T.S. and C.C.V. conceived the ideas. L.S., O.L., G.P. and C.C.V. provided materials and infrastructure. L.S., O.L., L.M., G.P and C.C.V. assisted in data collection. T.S. and S.R. quantified the behavioral responses. T.S. and L.S. conducted the statistical analyses and led the writing. All authors evaluated and approved the final draft of the article.

## Competing interests

The authors declare that they have no competing interests.

## Notes

### Competing Interest Statement

The authors have declared no competing interest.

