## Supplementary for "*In situ* novel environment assay reveals acoustic exploration as a repeatable behavioral response in migratory bats"

### Supplementary Information

#### Supplementary Tables

**Supplementary Table 1.** Descriptive statistics of response measures.

| Category | Response measure | Min | 1st Q | Median | Mean | 3rd Q | Max | N/A |
|---|---|---|---|---|---|---|---|---|
| Emergence | Body emergence latency (s) | 0.91 | 16.56 | 48.96 | 51.09 | 79.96 | 161.76 | 20 |
| | Head emergence latency (s) | 0.18 | 11.32 | 33.36 | 41.75 | 68.53 | 146.75 | 16 |
| | Emergence duration (s) | 0.56 | 3.16 | 4.92 | 10.61 | 10.14 | 114.12 | 20 |
| Spatial | Number of chambers visited | 1 | 3 | 9 | 9 | 12 | 25 | 20 |
| | Number of unique chambers visited | 1 | 3 | 6 | 5.25 | 7 | 9 | 20 |
| Acoustic | Number of calls after body emergence | 170 | 946 | 1227 | 1177 | 1482 | 2127 | 20 |
| | Number of calls after head emergence | 238 | 994 | 1274 | 1250 | 1574 | 2342 | 20 |
| | Number of calls during emergence | 4 | 24 | 56 | 72.81 | 100.5 | 335 | 20 |
| Other | Number of head pokes | 0 | 0 | 1 | 1.44 | 2 | 7 | 20 |
| | Number of air puffs | 0 | 3 | 5 | 5.75 | 8 | 22 | 20 |
| | Upside down crawling | Yes (n = 23) | | | No (n = 68) | | | 20 |

**Supplementary Table 2.** Spearman correlations between response measures within the same behavioral category.

| Category | Response measure | Response measure | $r_s$ | p |
|---|---|---|---|---|
| Emergence | Body emergence latency (s) | Head emergence latency (s) | 0.93 | < 0.001 |
| | Emergence duration (s) | Head emergence latency (s) | 0.25 | 0.017 |
| | Emergence duration (s) | Body emergence latency (s) | 0.51 | < 0.001 |
| Spatial | Number of chambers visited | Number of unique chambers visited | 0.89 | < 0.001 |
| Acoustic | Number of calls after body emergence | Number of calls after head emergence | 0.98 | < 0.001 |
| | Number of calls during emergence | Number of calls after head emergence | 0.34 | 0.001 |
| | Number of calls during emergence | Number of calls after body emergence | 0.22 | 0.04 |

**Supplementary Table 3.** Linear mixed model summary using Type II Wald Chi-square tests for effects of independent variables on 'Acoustic exploration' (residuals of the number of echolocation calls as function of the number of chambers visited). Individual ID was added as random effect. Continuous variables were scaled. Significant effects are reported in bold. Number of observations = 85.

| | Estimate | SE | $\chi^2$ | df | p |
|---|---|---|---|---|---|
| **Test (Second)** | **-344.60** | **52.43** | **43.20** | **1** | **< 0.001** |
| **Sex (Male)** | **144.71** | **69.44** | **4.34** | **1** | **0.04** |
| Weight* | 5.38 | 60.04 | 0.008 | 1 | 0.93 |
| Forearm | 22.30 | 38.74 | 0.33 | 1 | 0.56 |
| **Direction (North-South)** | **-93.99** | **43.81** | **4.60** | **1** | **0.03** |
| **Direction (East-West)** | **85.95** | **42.36** | **4.12** | **1** | **0.04** |
| Time since sunset* | -55.88 | 28.70 | 3.79 | 1 | 0.052 |
| Supplemental food (Yes)* | -113.33 | 71.10 | 2.54 | 1 | 0.11 |
| Orientation test (Before)* | 28.35 | 133.61 | 0.05 | 1 | 0.83 |

**VIF > 2.5, interpret results with caution*

**Supplementary Table 4.** Generalized linear mixed model summary using Type II Wald Chi-square tests for effects of independent variables on the number of chambers visited (family = Poisson). Individual ID was added as random effect. In addition, observation number was added to prevent overdispersion. Continuous variables were scaled. Number of observations = 85.

| | Estimate | SE | $\chi^2$ | df | p |
|---|---|---|---|---|---|
| Test (Second) | -0.28 | 0.14 | 3.75 | 1 | 0.053 |
| Sex (Male) | 0.04 | 0.22 | 0.03 | 1 | 0.86 |
| Weight* | -0.07 | 0.14 | 0.26 | 1 | 0.61 |
| Forearm | -0.03 | 0.11 | 0.11 | 1 | 0.74 |
| Direction (North-South) | -0.03 | 0.12 | 0.08 | 1 | 0.77 |
| Direction (East-West) | 0.01 | 0.12 | 0.01 | 1 | 0.91 |
| Time since sunset* | 0.05 | 0.08 | 0.41 | 1 | 0.52 |
| Supplemental food (Yes)* | -0.24 | 0.18 | 1.81 | 1 | 0.18 |
| Orientation test (Before)* | 0.29 | 0.36 | 0.64 | 1 | 0.42 |

**VIF > 2.5, interpret results with caution*

**Supplementary Table 5.** Linear mixed model summary using Type II Wald Chi-square tests for effects of independent variables on the body emergence latency (log-transformed). Individual ID was added as random effect. Continuous variables were scaled. Significant effects are reported in bold. Number of observations = 104.

| | Estimate | SE | $\chi^2$ | df | p |
|---|---|---|---|---|---|
| Test (Second) | -0.34 | 0.23 | 2.31 | 1 | 0.13 |
| Sex (Male) | -0.003 | 0.31 | 0.0001 | 1 | 0.99 |
| **Weight*** | **0.38** | **0.19** | **4.05** | **1** | **0.04** |
| Forearm | 0.07 | 0.15 | 0.23 | 1 | 0.63 |
| Direction (North-South) | 0.23 | 0.16 | 2.22 | 1 | 0.14 |
| Direction (East-West) | 0.07 | 0.16 | 0.19 | 1 | 0.66 |
| Time since sunset* | 0.09 | 0.10 | 0.79 | 1 | 0.37 |
| **Supplemental food (Yes)*** | **1.31** | **0.34** | **14.57** | **1** | **< 0.001** |
| Orientation test (Before)* | -0.08 | 0.54 | 0.02 | 1 | 0.88 |

**VIF > 2.5, interpret results with caution*

**Supplementary Table 6.** Linear mixed model summary using Type II Wald Chi-square tests for effects of response measures exclusively added as independent variable to the final model for 'Acoustic exploration' (residuals of the number of echolocation calls as function of the number of chambers visited). Individual ID was added as random effect. Significant effects are reported in bold. Number of observations = 85.

| | Estimate | SE | $\chi^2$ | df | p |
|---|---|---|---|---|---|
| Body emergence latency (s, log-transformed) | 4.70 | 26.85 | 0.03 | 1 | 0.86 |
| **Residuals (calls ~ emergence duration)** | **2.36** | **0.57** | **17.18** | **1** | **< 0.001** |
| **Head pokes in adjacent rooms** | **57.87** | **19.16** | **9.12** | **1** | **0.003** |
| Number of air puffs | 7.71 | 7.85 | 0.96 | 1 | 0.33 |
| Upside down (Yes) | -128.15 | 68.94 | 3.46 | 1 | 0.06 |

#### Supplementary Figures

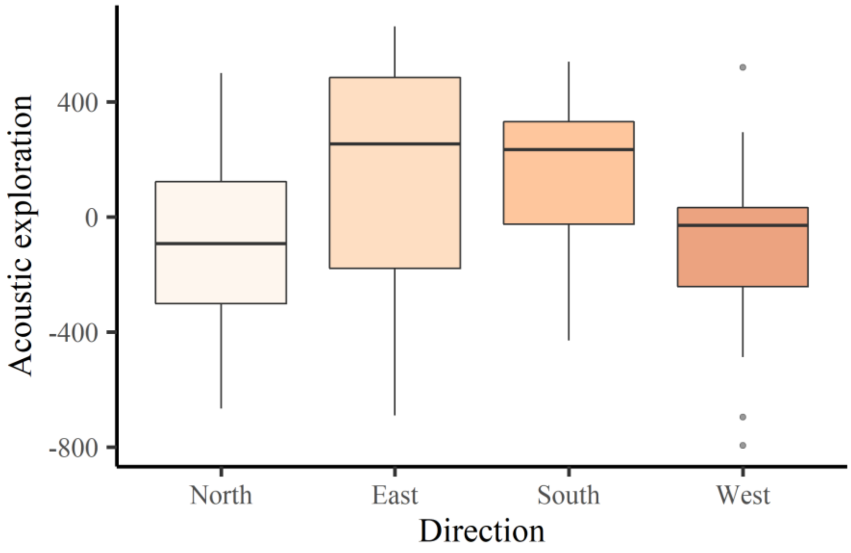


**Supplementary Figure 1. Acoustic exploration (number of echolocation calls relative to the number of chambers visited) as function of emergence direction.** Note that South-East is the local migratory direction.
